# The molecular architecture of the kidney slit diaphragm

**DOI:** 10.1101/2023.10.27.564405

**Authors:** Alexandra N. Birtasu, Konstantin Wieland, Utz H. Ermel, Johanna V. Rahm, Margot P. Scheffer, Benjamin Flottmann, Mike Heilemann, Florian Grahammer, Achilleas S. Frangakis

## Abstract

Vertebrate life depends on renal function to filter excess fluid and remove low-molecular-weight waste products. An essential component of the kidney filtration barrier is the slit diaphragm (SD), a specialized cell-cell junction between podocytes. Although the constituents of the SD are largely known, its molecular organization remains elusive. Here, we use super-resolution correlative light and electron microscopy to quantify a linear rate of reduction in albumin concentration across the filtration barrier. Next, we use cryo-electron tomography of vitreous lamellae from high-pressure frozen native glomeruli to analyze the molecular architecture of the SD. The resulting densities resemble a fishnet pattern. Fitting of Nephrin and Neph1, the main constituents of the SD, results in a complex interaction pattern with multiple contact sites between the molecules. Using molecular dynamics flexible fitting, we construct a blueprint of the SD, where we describe all interactions. Our architectural understanding of the SD reconciles previous findings and provides a mechanistic framework for the development of novel therapies to treat kidney dysfunction.

## Introduction

In vertebrates, kidneys are critical in maintaining and regulating water and electrolyte homeostasis. They clear low-molecular-weight waste products from large volumes of plasma while retaining proteins in the bloodstream. The kidney’s filtration system, the glomerular filtration barrier, is located within microvascular units called glomeruli, which consist of three layers: a monolayer of fenestrated endothelial cells; a thick glomerular basement membrane (GBM); and highly specialized, terminally differentiated epithelial cells called podocytes ^12^. Protrusions of the podocytes, known as foot processes, interdigitate with those of adjacent podocytes and create a unique cell-cell contact called the slit diaphragm (SD) that is composed of Nephrin and Nephrin-like 1 (Neph1)^3^. Proper filter and podocyte function is dependent on the integrity of the SD. Many kidney diseases are associated with SD dysfunction and pathological changes in the SD have been described in acquired and hereditary diseases that lead to proteinuria, such as minimal change disease, congenital nephrotic syndrome and nephrotic syndrome or diabetes ^45^. Proteinuria has for long been established as an independent renal and cardiovascular health risk factor ^6^, which emphasizes the importance in understanding the SD structure and function at the molecular level to foster development of antiproteinuric therapies.

The SD has been studied extensively using a variety of techniques. Early electron micrographs suggested a zipper-like structure for the SD that would imply a filter function and facilitate size selectivity ^7,8^. Cryo-electron microscopy of vitreous sections showed individual densities spanning the gap between podocytes, suggesting that the repetitive Ig-like C2-type domains (hereafter abbreviated to Ig*X,* with X=1,2,..,9 indicating the number of the Ig domain) of Nephrin, Neph1 and podocin provide the SD with inherent flexibility ^9^. Proteomics studies have revealed that the core proteins Nephrin, Neph1 and podocin form a supramolecular SD complex ^10^. Super-resolution microscopy has been used to pinpoint the location of the core proteins and suggest their stoichiometry ^11,12^. X-ray crystallography was utilized to solve the structure of the complex comprising SYG-1 and SYG-2, homologs of Neph1 and Nephrin in *Caenorhabditis elegans*, respectively, which displays heterophilic *trans* interactions ^13^. Advanced light microscopy and electron microscopy with tracers have been used to investigate the selectivity properties of the filter under physiological conditions by visualizing the distribution of proteins at the filtration barrier ^14^. Detailed physical models have attributed the size and charge selectivity of the glomerular filtration barrier to streaming potentials, which prevent most of the filtrate from entering the filtration barrier at the endothelial cells ^15^. The function of the SD has been emphasized by its role in providing mechanical resistance against blood pressure ^12^. Despite this wealth of information, the molecular architecture of the SD remains elusive due to the challenge of preparing tissue samples for electron microscopy, the difficulty in obtaining high-resolution images of flexible molecules, and the introduction of fixation artifacts as a result of insufficient preservation methods.

Here we use cryo-focused-ion-beam (FIB)-milled high-pressure frozen glomeruli and cryo-electron tomography (cryo-ET) to visualize the SD at a resolution sufficient to discern the scaffold of individual Nephrin and Neph1 molecules. To verify the role of the SD in the filtration mechanism, we measure the precise distribution of fluorescently labelled albumin at the glomerular filtration barrier with correlative super-resolution fluorescence and electron microscopy. Subtomogram averaging and molecular dynamics flexible fitting allow us to map heterophilic and homophilic interactions between Ig domains from Nephrin and Neph1 to reveal the overall architecture of the SD at molecular resolution.

## Results

### A substantial amount of albumin passes through the SD under no-flow conditions

The current electrokinetic model of renal filtration postulates that under physiologic conditions albumin molecules never reach the SD ^16,12 15^. Only under pathological conditions, such as podocyte diseases or conditions in which capillary flow ceases, do albumin molecules traverse the GBM and the SD. To probe the electrokinetic model and understand whether the SD is involved in active filtration, we analysed the static filtration properties of the SD by correlative super-resolution light and electron microscopy under capillary no-flow conditions. Fluorescently labelled albumin (Alexa Fluor^TM^ 555 conjugated to BSA, approximately 66 kDa) was injected into the kidney artery of mice (Extended Data Figure 1). After 60 seconds, microbiopsies (n=5) were collected, fast high-pressure frozen, freeze-substituted and ultrathin sectioned. We used direct stochastic optical reconstruction microscopy (dSTORM) to image albumin in the thin sections ^17^ and then recorded large landscapes of the glomerular section in the electron microscope, showing all characteristic landmarks (Figure 1a). The obtained images were registered with affine transformations and superimposed to account for the different information present in the microscopy methods. This allowed us to assign fluorescent clusters of albumins to individual compartments of the glomerulus (Figure 1a,b). The mean concentration of albumin clusters was highest in the lumen of the capillaries (2.2 ± 1.0 clusters/µm^2^, n=28), lower in the GBM (1.6 ± 0.8 clusters/µm^2^, n=28), and lowest in the lumen of the urinary space (1.0 ± 0.9 clusters/µm^2^, n=28) (Figure 1c). One would anticipate that most of the albumin is held back by the filtration barrier, thus most of the albumin should be found in the LC and nearly nothing in the LUS. However, we still observe a substantial amount of albumin in LUS even though the SD seems intact in the electron micrograph.

**Figure 1:**
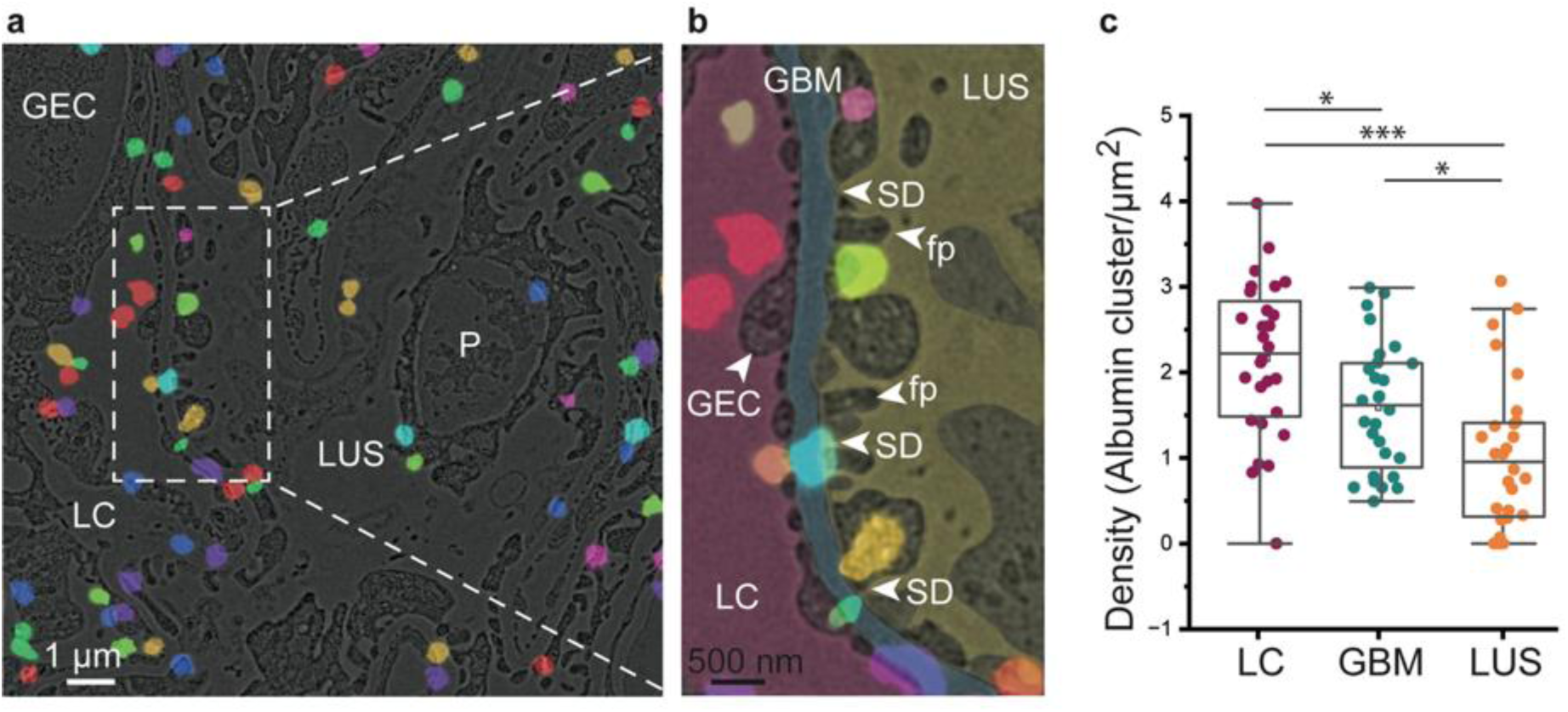
Super-resolution correlative light and electron microscopy displays the positions of albumin in the vicinity of the slit diaphragm. **a,** Electron micrograph of a high-pressure frozen and freeze-substituted kidney section showing the luminal capillary (LC), glomerular endothelial cells (GEC), podocyte (P), and the lumen of the urinary space (LUS). The positions of fluorescent albumin clusters are shown in different colors. Albumin was labeled with Alexa Fluor 555. **b,** Close-up of the micrograph shown in (a) with the manual segmentation of the areas of the LC (magenta) covered by GEC (light magenta), the glomerular basement membrane (GBM, light blue), and the podocyte with characteristic foot processes (fp and SD indicated by white arrowhead) in the LUS (yellow). **c,** Statistical evaluation of the albumin density in the glomerular filtration barrier measured in albumin clusters per µm^2^ (for n=28). A linear reduction in the concentration can be observed. Paired t-test: LC v LUS p=5.3×10^-6^, GBM v LC p=0.0033, GBM v LUS p=4.56×10^-4^.

### Cryo-electron tomograms of vitreous lamellae display the scaffold of individual Nephrin and Neph1 molecules

To overcome the limitations of plastic-embedded samples and gain insight into the molecular architecture of the native renal glomerular filtration barrier, we performed cryo-electron tomography on lamellae of high-pressure frozen, freshly isolated mouse kidney glomeruli (Extended Data Figure 2). First, we localized the glomeruli using cryo-confocal laser scanning microscopy (Figure 2a) and labelled their positions for cryo-focused ion beam (FIB) milling. Second, we cryo-FIB-milled 130–250 nm thick on-grid lamellae, which were then imaged in the transmission electron microscope (Figure 2b). Due to the thickness of the frozen sample (∼40 µm), the lamellae had to be cut at a steep angle (∼45 degrees). This is extremely tedious to mill (∼100 times more material needs to be removed compared to lamellae from a plunge-frozen eukaryotic cells) and reduces the effective angular range of the subsequent tilt-series acquisition, leading to tomograms with a large missing wedge (∼100 degrees, Extended Data Movie 1). In its native frozen-hydrated state, all characteristic landmarks of the glomerular tissue organization can be seen, but at higher resolution and better preservation compared to the plastic embedded samples (Figure 2c,d,e, Extended Data Movie 2). The foot processes are filled with actin bundles that are branching in the vicinity of the SD (Figure 2d,e, Extended Data Movie 1 and 2).

**Figure 2:**
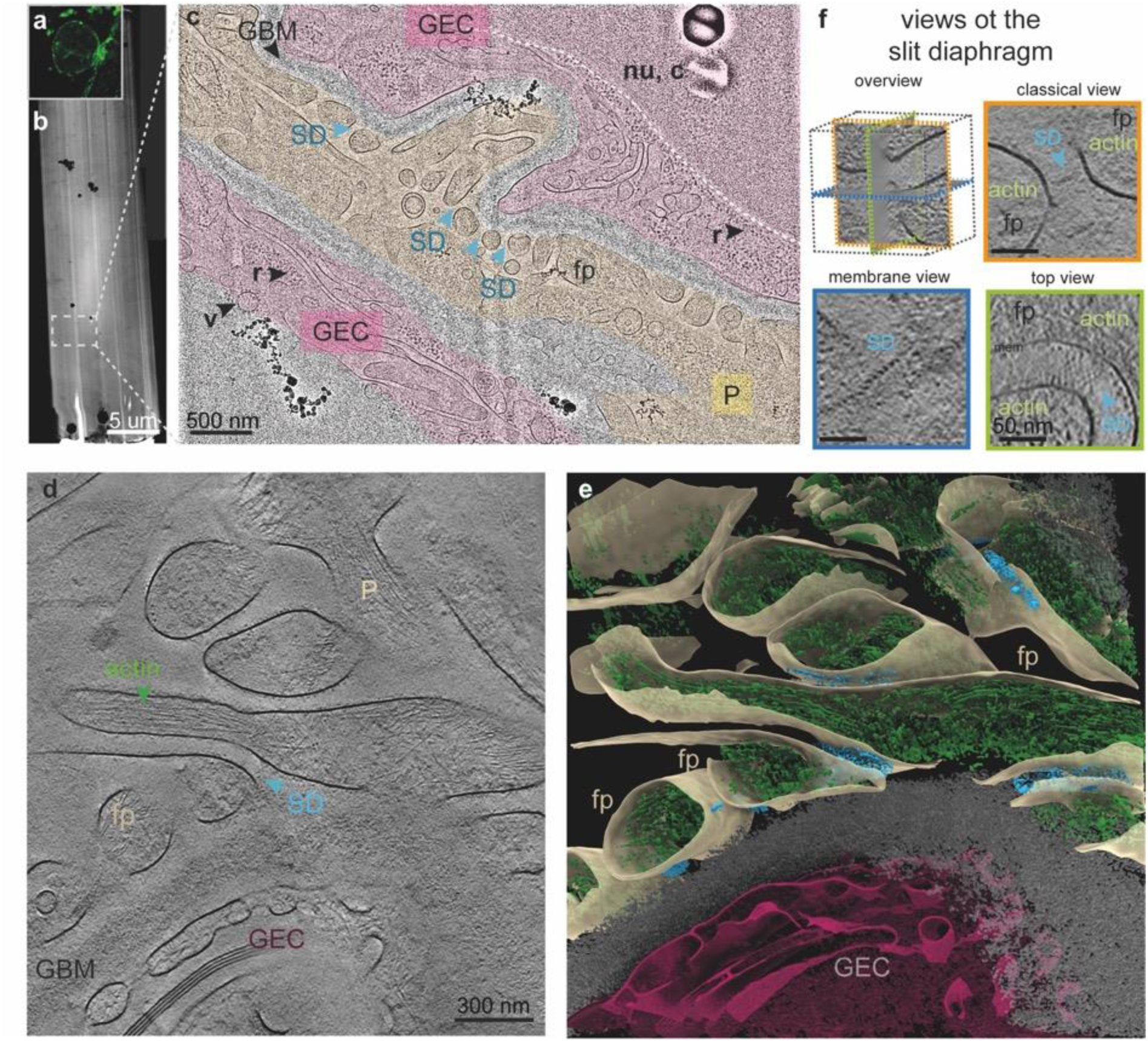
Visualization of the murine kidney slit diaphragm from high-pressure frozen, cryo-focused ion beam milled lamella of a native mice glomerulus. **a,** A low-magnification image of a vitreous lamella through the kidney tissue. **b,** Fluorescent image of the glomerulus from murine kidney tissue from which the lamella was produced. The glomerulus appears as a roundish object with a diameter of ∼140 µm. **c,** Magnified image of the region of interest where a tomogram was recorded. All the characteristics and landmarks of the glomeruli morphology can be seen. The layered organization of the filtration system is shown: A glomerular endothelial cell (GEC, magenta) with its nucleus (nu), ribosomes (r) in the cytoplasm (c), and the glomerular basement membrane (GBM, grey) with the foot processes (fp) of the podocytes (P, yellow). In between the foot processes, the SD can be discerned (light blue arrows). **d,** 1 nm thick slice through the cryo-electron tomographic reconstruction of the filtration system. The foot processes (fp, yellow) of the podocytes (P) are filled with actin bundles (green). Near the SD, actin appears more branched and points towards the membranes. The fenestrations of the GEC (magenta) are directly next to the GBM (grey). The SD (light blue arrows) can also be seen in arbitrary views. **e,** Isosurface representation of the whole tomogram shown in (d). Foot processes (fp, yellow) are filled with actin bundles (green) with the densities at the area of the SD (blue). The fenestrations of the GEC cover the GBM (grey). **f,** Three orthogonal views through the SD; the colors of the borders correspond to the orthogonal slices in the overview. The classical view, as also seen in conventional electron microscopy images, is shown with an orange border. The membrane view (with a dark blue border) displays the cross-sections through individual strands (either Nephrin or Neph1). The top view (with a dark green border) displays molecular strands intertwining and spanning two foot processes (fp) filled with actin. The curvature of the foot processes is evident.

Importantly, the SD can be seen from different viewpoints (Figure 2f), revealing a picture that is much more differentiated than those obtained previously ^9^. The (classic) side view appears similar to that of previous studies (Figure 2f). The SD has a width of ∼53.1 nm (±3.2 nm, n=51; Figure 2f, classical view) in the extracellular space between two adjacent foot processes, which is slightly larger than previously described^9^. This discrepancy can be explained by the absence of sample shrinkage or compression in the native vitreous lamellae, unlike in other sample preparation methods. The top view displays molecular strands that form an apparent fishnet pattern (Figure 2f, top view) and have a distinct lateral spacing of ∼12.3 nm (±1.6 nm, n=150), spanning the gap between adjacent foot processes. They can be seen extending from one side of the membrane for ∼15 nm and then bending ∼40 degrees to reach the opposite foot process. Two layers of crossing strands can be discerned. In the membrane view, the individual strands can be seen in a cross-section (Figure 2f, membrane view).

### The sub-tomogram average shows a quasi-periodic fishnet pattern

To further investigate the structural arrangement of the extracellular parts of the proteins forming the SD, we performed sub-tomogram averaging. From 47 tilt series, we selected those with the best alignment and thinnest lamella (<180 nm), resulting in six cryo-electron tomograms with 43 segments of the SD for further analysis (Extended Data Table 1). Sub-tomogram averaging and subsequent refinement yielded in a final set of 134 sub-tomograms, each of which contained approximately ten molecular strands. The sub-tomogram average (Figure 3a, Extended Data Figure 3) shows a quasi-periodic arrangement of the molecular strands, as well as the fishnet pattern that is visible in the reconstructions (Figure 2f). Despite the high flexibility of the molecules, the varying intermembrane distance (Figure 3c) and the curved nature of the podocytes, a long-range periodicity can be observed (Figure 3a, top and membrane views, Extended Data Figure 3c). As views of the SD from all directions contributed to the average, the resulting cryo-ET density map is not affected by the large missing wedge of the reconstruction and depicts all SD arrangements seen in the tomograms (Figure 2f, Figure 3a, Extended Data Figure 3b).

**Figure 3:**
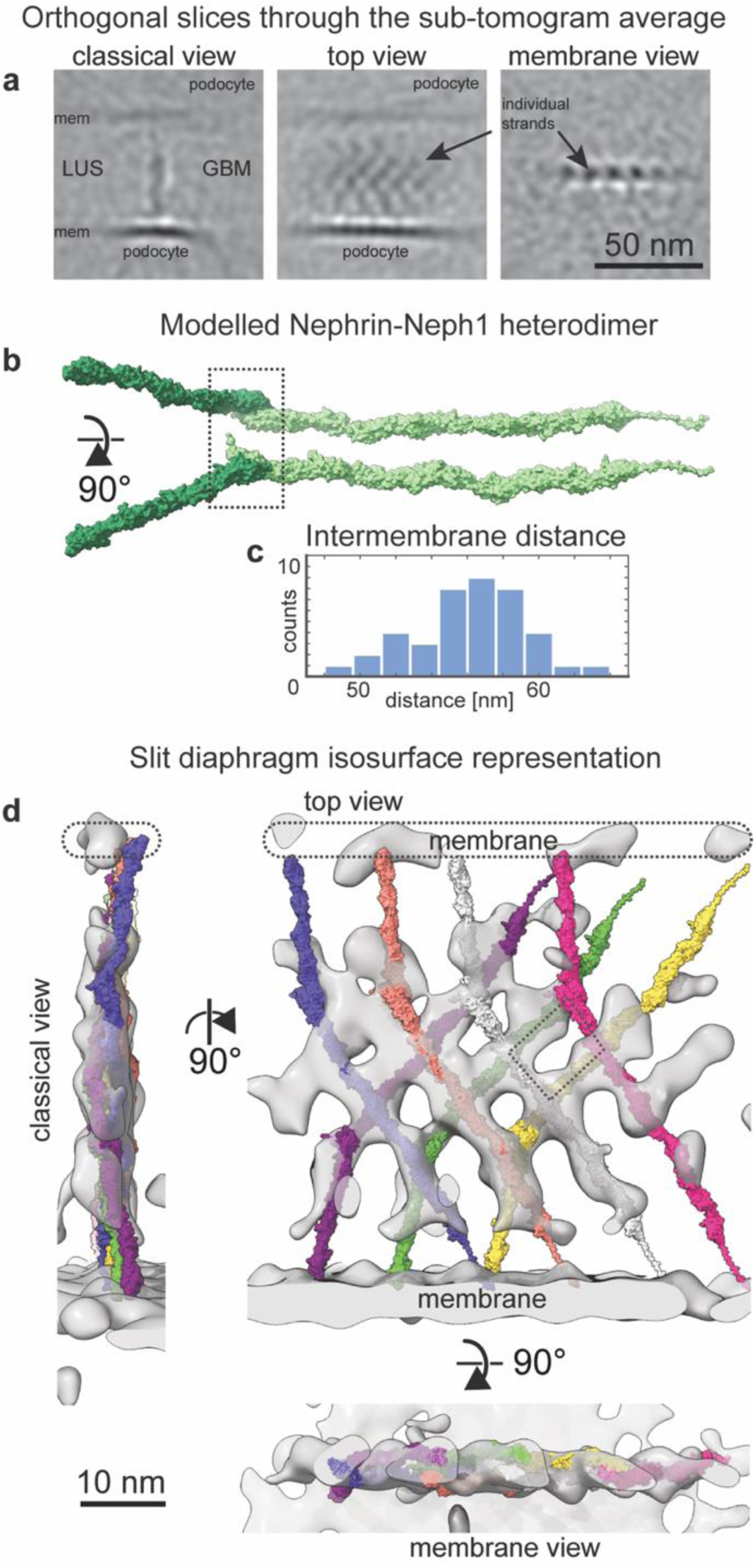
The SD is a network of Neph1 and Nephrin, creating a fishnet pattern. **a,** Three views of 0.4 nm thick computational sections through the sub-tomogram average. They resemble the individual views seen in the reconstructions (Figure 2) with an improved signal-to-noise ratio. The first membrane appears prominently, while the juxtaposed membrane is being smeared out. **b,** Two views of the Nephrin-Neph1 (ID: AF-Q9QZS7-F1, AF-Q80W68-F1) heterodimer as modelled with AlphaFold ^18,19^ and based on the interface of the SYG-2-SYG-1 crystal structure (PDB: 4OFY). **c,** Intermembrane distances of the SD as measured from the sub-tomogram average. The large variation is due to the pleiomorphic shape of the SD. **d,** Three views of the Nephrin-Neph1 heterodimer (each shown in different colors to maximize the contrast) rigidly fitted into the sub-tomogram average (shown in transparent grey). An individual heterodimer spans the distance between the two membranes. A fishnet pattern is created by the densities of individual heterodimers crossing over each other. Four heterodimers form a rhomboid shape (indicated by the dotted rhomboid shape) at the center of the SD.

### Nephrin and Neph1 heterodimers sandwich and periodically repeat one-dimensionally to form the SD

Structural comparison of Nephrin and Neph1 (which engage in the formation of the extracellular part of the SD) with homologs SYG-2 and SYG-1, suggests a heterophilic binding mode for their Ig1 domains ^13^ (Extended Data Figure 4a). Thus, we generated molecular models of Nephrin and Neph1 using AlphaFold (IDs: AF-Q9QZS7-F1, AF-Q80W68-F1). We assembled these models into a heterodimer based on the interface of the crystal structure of SYG-2-SYG-1 (PDB: 4OFY) by aligning the Ig1 domains of Nephrin and Neph1 to those of SYG-2 and SYG-1, respectively (Figure 3b, Extended Data Figure 4b). The complete atomic model of the heterodimer appears as an elongated molecular strand with an angle of ∼150 degrees between Nephrin and Neph1. The model of the heterodimer was then rigid-body fitted into the cryo-ET density (Figure 3d, Extended Data Movie 3). Progressive coarse fitting of the cryo-ET density with the heterodimers resulted in a complete coverage of the densities of the SD. The resulting arrangement of the heterodimers indicates several homophilic and heterophilic interactions and provides numerous insights. The ratio of Nephrin to Neph1 is 1:1. The Nephrin molecule extends from the adjacent foot process at an angle of ∼50 degrees, while the Neph1 molecule extends from the opposing foot process at an angle of ∼90 degrees. The opposing heterodimer is rotated 180° around the normal vector of the classical view. Through this arrangement a symmetry is created, with immediate implications: Nephrin and Neph1 emanate from each podocyte and form a heterophilic *cis* interaction first and then heterophilic and homophilic *trans* interactions at the center of the SD. Indeed, podocytes and nephrocytes are the only two cell-types that express both Nephrin and Neph1 ^13^.

### The SD proteins Nephrin and Neph1 intertwine into a stable fishnet pattern

Next, we produced a quasi-crystalline density map from the cryo-ET density map by translational and rotational symmetrization (12.3 nm in direction of the normal vector to the classical view and 180 degrees around the normal vector to the classical view) (Figure 4). The fit of the heterodimers in the symmetrized density map was then improved by molecular dynamics flexible fitting (MDFF) (ChimeraX-Isolde ^20^). Distance restraints were applied separately for each Ig domain and for the glutamines mediating the interaction interface in the Ig1 domains (Extended Data Figure 4), but not for the linker regions between them. In the resulting arrangement, several interactions between the Nephrin-Neph1 heterodimer and other Nephrin-Neph1 molecules can be observed (Figure 4a, Extended Data Movie 4):

i. A heterophilic *cis* interaction (from the same cell) between Neph1-Ig2 and Nephrin-Ig8/Ig9 (Figure 4b)
ii. A homophilic *trans* interaction between Nephrin-Ig2/Ig3 and Nephrin-Ig5 (Figure 4c)
iii. A homophilic *trans* interaction between Nephrin-Ig5 and Nephrin-Ig2/Ig3 (Figure 4c)
iv. A heterophilic *cis* interaction between Nephrin-Ig8/Ig9 and Neph1-Ig2 (Figure 4b).

**Figure 4:**
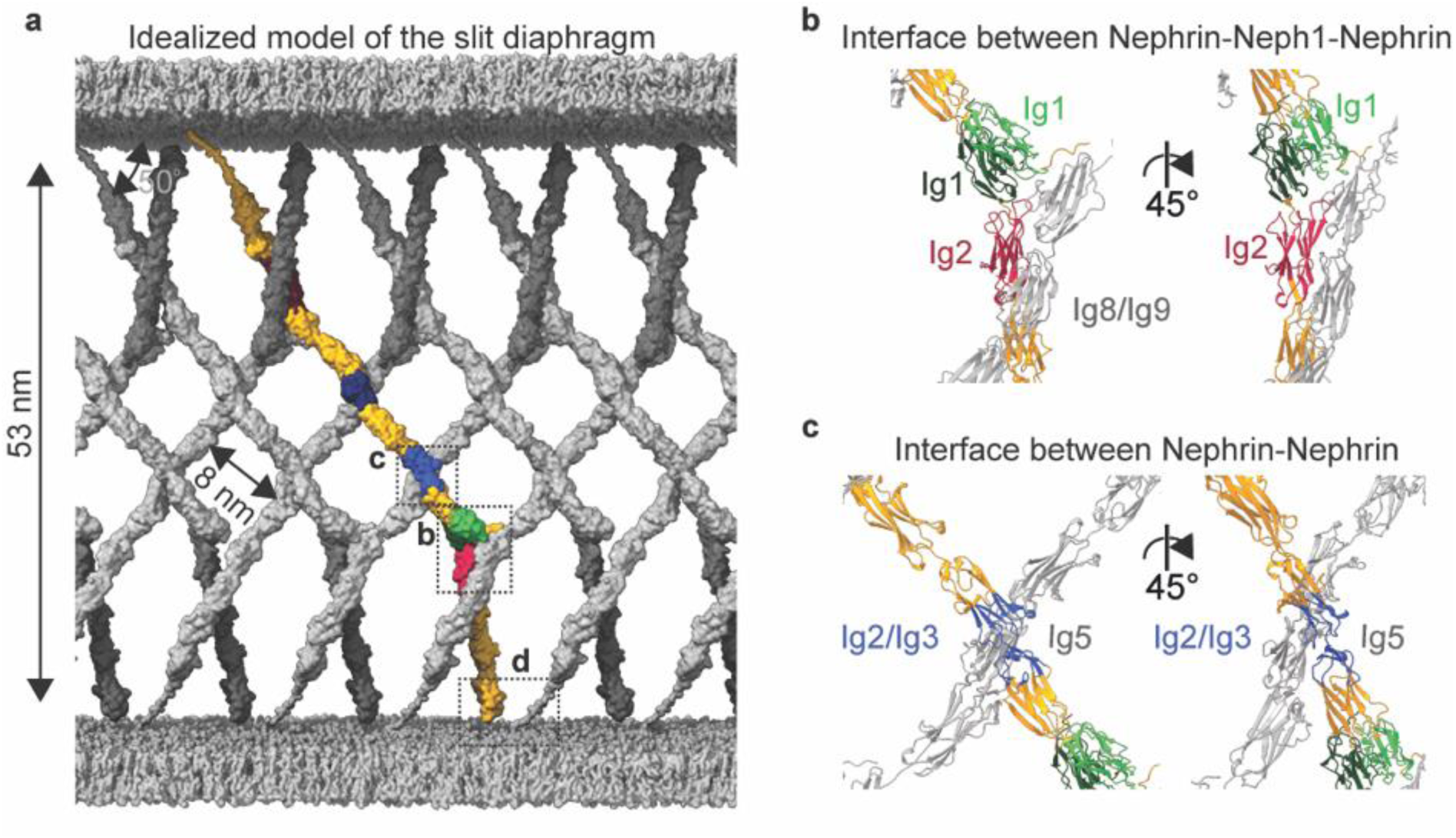
Idealized model of the slit diaphragm. **a,** Idealized model of the SD after molecular dynamics flexible fitting and using the Nephrin-Neph1 heterodimer as a unit-cell in one-dimensional crystal packing. One heterodimer is highlighted in color (gold, the positions where individual extracellular domains (Ig*X*) are predicted to interact are shown in a different color), while all other heterodimers are shown in grey. Each heterodimer is expected to interact with (at least) four other heterodimers ^23^. **b,** Heterophilic *cis* interface between Nephrin-Ig8/Ig9 (grey) and Neph1-Ig2 (wine red) emanating from the same membrane. The *trans* interface between Nephrin-Ig1 and Neph1-Ig1 is shown in green. The interface between Nephrin-Ig8/9 and Neph1-Ig2 is shown in wine red. One heterodimer is involved with the same interface two times. **c,** Homophilic *trans* interaction between two Nephrin molecules via their Ig2/Ig3 (light blue) and Ig5 (grey) domains. The Nephrin molecules create an additional homophilic interaction, as in (c), between their Ig5 (dark blue) and Ig2/3 (grey) domains. **d,** The close proximity of the cytoplasmatic tails of Nephrin and Neph1 would allow for intracellular interactions.

The orientation of the heterodimer within the SD is determined by the Nephrin-Ig1-Neph1-Ig1 interface, resulting in the distinct “fishnet” pattern. Analysis of the individual domain interfaces reveals that the opposing electrostatic charges within Nephrin Ig2/Ig3 and Nephrin Ig5 facilitate robust attachment (Extended Data Figure 5). In addition, asparagine (N)-linked glycosylation sites (Uniprot accession: Q9QZS7) have been predicted within the Nephrin Ig1, Ig4 and Ig6 domains, which could be modified by different glycans ^21^. These glycans serve as impediments for interactions and effectively hinder lateral domain mobility (Extended Data Figure 6, Extended Data Movie 5). Notably, this glycosylation-mediated obstruction is also evident in the context of the glycan on Neph1-Ig1 (Uniprot accession: Q80W68), which is oriented towards the terminus next to Nephrin-Ig2.

It is therefore striking that the interaction interfaces repeat. The *cis* interaction between Nephrin and Neph1 can manifest either as a domain interaction or as a more extended interaction involving several subunits. Thus, each Nephrin-Neph1 heterodimer experiences at least four interactions with other heterodimers (additional interactions may occur in the vicinity of the plasma membrane), in addition to the heterophilic *trans* interaction between the Nephrin and Neph1 molecules in the heterodimer itself. This architecture creates a sandwich arrangement with almost regularly spaced Nephrin-Neph1 heterodimers that can only extend in the direction orthogonal to the classical view, which allows for the creation of a diaphragm. This contrasts with other junctions that create a quasi-two-dimensional crystals, which can be extended infinitely in two dimensions ^22^.

Architecturally, this model includes structural holes on the order of 7×7 nm^2^ and larger, which would allow the diffusion of albumin into the urinary space while potentially prohibiting larger macromolecular complexes from passing through. Considering the pleiomorphic shape of the podocytes as well as the various shear forces exerted on the SD, this multitude of interactions creates a much more stable arrangement than of an individual protein complex spanning the two podocytes, without additional interactions. Therefore, between the individual Ig domains, a stable and robust network can be built.

## Discussion

As failure of the SD causes proteinuria and kidney disease, the SD must remain intact and stable for long periods of time ^4^ ^5^. Our cryo-ET density reveals that each Nephrin-Neph1 heterodimer is interacting with at least four other heterodimers, which together create a fishnet pattern that gives this specialized cell-cell junction both high structural stability and flexibility. Near the podocyte plasma membrane additional interactions between Nephrin and Neph1 are also possible ^24^. The fishnet pattern of the SD allows for changes in the distance between adjacent podocytes, without affecting their points of interaction. The distance compensation happens via a change in the angles of the rhomboid shape formed between the heterodimers, while the side length remains the same. In this model, the linker regions between the Ig domains may act as pivot points, enabling the SD to change its width without needing to unfold the Ig domains. In this way, changes in blood pressure within glomerular capillaries can be compensated for smoothly by the SD. This is needed to filter 180 litres of blood on a daily basis, over an average human lifespan of eight decades ^25^.

Our model shows that the center of the SD is composed of Nephrin molecules alone, which are solely responsible for the *trans* interactions between the podocytes and thus Nephrin has a more important role for the slit maintenance. This is corroborated by studies in animal models using either siRNA-mediated knock-down or inducible deletion of the *Neph1* gene, in which Nephrin maintains the structured SD for a long time before albuminuria occurs ^10,26^. On the other side knock-downs of *NPHS*1 result into massive proteinuria and neonatal death ^27^. Furthermore, our model is consistent both with the compression model ^16,12^, in which the SD is required for a stable podocyte network that compresses the GBM to potentially reduce permeation speed, and with the electrokinetic model ^15^, in which negatively charged molecules are moved back to the capillaries by electrophoresis, which actively prevents the clogging of the glomerular filtration barrier. Our model suggests that the ratio of Nephrin to Neph1 is 1:1, which differs from proteomics studies that suggest a ratio of 1.6:1 ^10^. However, Nephrin and Neph1 was analyzed in all podocyte membranes, so their findings cannot be restricted to the SD alone.

Several known disease-causing NPHS1 mutations disrupt either protein translation or translocation to the plasma membrane ^5,28 29^. In these cases, formation of the SD is not possible, thereby resulting in protein urea and renal failure. When the SD is formed, its fishnet architecture inherently affords a wide spectrum of mechanical stability and flexibility through multiple interactions between heterodimers, enabling the SD to withstand varying degrees of positive and negative pressure (for example, under no-flow conditions). Elucidation of the molecular structure of the SD will help to clarify pathogenic phenotypes of SD malformation at a molecular level. Autoimmune responses that directly target the SD, as in the case of glomerulonephritis after kidney transplantation, can now be understood since the sites of SD accessibility are revealed^30^.

## Data Availability Statement

The cryo-ET structure solved in this study is available in the Electron Microscopy Data Bank (EMDB) under the accession codes EMD-XXXXX. The model is available under the accession codes EMD-XXXXX.

## Acknowledgements and Funding

We thank the Frankfurt Center for Electron Microscopy and the Frankfurt Center for Advanced Light Microscopy for measurement time. We thank C.J.O. Kaiser for introducing us to the high-pressure-freezing method and providing on-site assistance, N. Alivodej for providing us with mice kidneys, and D. Bublak for assistance with formvar-coating. A. Habermann for assistance during plastic embedding. We thank T. Sewell, S.M. Arghittu and R. Covino for valuable discussions and comments about MDFF. We thank L. Schaefer for the glomeruli isolation protocol. FG was supported by DFG (CRC 1192, GR3933/1-1). A.N.B., J.V.R. and M.H. were funded by Research Training Group iMOL (GRK 2566/1). A.S.F. was supported by the Deutsche Forschungsgemeinschaft (FR 1653/14-1 for U.H.E., and FR 1653/14-1 for K.W.).

## Author Contributions

A.N.B: Planned and carried out glomeruli preparation, recorded and processed tomography datasets. A.N.B. and K.W.: Performed high-pressure-freezing and acquired cryo-LSM images of frozen glomeruli. K.W.: Planned and carried out cryo-FIB sample preparation. U.H.E: Provided tomography analysis pipelines and assisted during processing. B.F. and M.H.: Planned and carried out super-resolution microscopy experiments. J.V.R.: Processed dSTORM data sets of kidney sections. J.V.R. and A.N.B.: Analyzed albumin clusters of kidney sections. A.N.B., M.P.S. and A.S.F.: Built the models. F.G.: Supervised research and carried out mouse experiments. A.S.F.: Designed and supervised research. A.N.B. and A.S.F.: Wrote the manuscript, with contributions from all authors.

## Materials & Methods

The research presented in herein complies with all relevant EU, national and regional laws and regulations. We also comply with EU, national and international ethics-related rules and professional codes of conduct.

### Animals for CLEM albumin localization

All experiments were conducted in accordance with the German animal welfare guidelines and the NIH Guide for the Care and Use of Laboratory Animals and were approved by the responsible authority (G-07/13 Regierungspräsidium Freiburg, Germany). C57Bl/6 mice were housed in a specific-pathogen-free (SPF) facility with free access to chow and water and a 12 h day/night cycle.

### CLEM albumin sample preparation – tissue processing

Animals were intraperitoneally injected with ketamine/xylazine (7 µL/kgBW, 7 g/kgBW). Kidneys were injected intravenously through the retroorbital plexus with 500 µg bovine serum albumin (BSA) (Alexa Fluor^TM^ 555 conjugate, Thermo Fisher Scientific Inc., Waltham, MA, USA). After 60 s, kidneys were dissected together with the abdominal aorta. The tissue was cut into small pieces and placed in gold-plated copper specimen carriers with 0.15 μm recess (type 665; Wohlwend, Sennwald, Switzerland) filled with 20% dextran. The tissue sections were high-pressure frozen in an HPM-010 (Abra Fluid, Widnau, Switzerland). Freeze substitution was performed in an automated system (EM AFS2, Leica Microsystems, Wetzlar, Germany) with a solution of 2% uranyl acetate (Serva, Heidelberg, Germany), 8% methanol and 2% water dissolved in glass-distilled acetone (Electron Microscopy Sciences, Hatfield, USA) at –90°C for 64 h and at –45°C for 24 h, followed by three washes in acetone at –45°C for 10 min. Tissue sections were sequentially infiltrated with increasing concentrations (10%, 25%, 50% and 75%) of Lowicryl HM20 (Polysciences, Hirschberg an der Bergstrasse, Germany) for 4 h each, while raising the temperature to –25°C in increments of 5°C. Lowicryl was exchanged two times after 12 h and a third time after 48 h. The tissue sections were then UV polymerized for 48 h at –25°C, followed by another 9 h at 20°C. Carriers were observed in a confocal microscope (LSM700, Carl Zeiss, Jena, Germany) to identify fluorescent glomeruli. Distances of glomeruli to tissue edges and/or carrier edges were measured. Ultrathin sections (40 nm) were cut with an ultramicrotome (Ultracut UC7, Leica Microsystems) and transferred to formvar-coated copper finder grids (Plano).

### Super-resolution imaging

Samples were imaged in PBS (Polyscience) with 50 – 200 mM MEA (Sigma-Aldrich, St. Louis, Missouri, US, #M6500-25G) added to adjust single-molecule photoswitching of the fluorophore ^31^ The average localization precision was 47 nm. The average localization precision was 47 nm. Measurements were conducted on a home-built microscope for single-molecule localization microscopy. In brief, an inverted microscope (IX81, Olympus, Tokyo, Japan) was equipped with a 150× UApo TIRFM oil objective lens (NA 1.45, Olympus) and operated in total internal reflections fluorescence (TIRF) mode. The fluorescence signal was collected on an Andor Ixon Ultra EMCCD chip (DU-897U-CS0-#BV; Andor Technology Ltd, Belfast, Northern Ireland). Alexa555 BSA Albumin (Thermo Fisher Scientific Inc.) was excited with a 561 nm diode laser (Cube; Coherent, Newton, CT, USA). Illumination intensities were adjusted to 0.65 kW/cm^2^ for direct stochastic optical reconstruction microscopy (dSTORM) imaging. We collected 3000 frames with an integration time of 200 ms, an image size of 512×512 px² with a pixel size of 108 nm, and an EM gain of 200.

### Localization and clustering of dSTORM movies

The dSTORM movies were processed using Picasso ^32^ (version 0.4.11) to obtain single-molecule localizations. In a first step, localizations were found by using the integrated Gaussian maximum likelihood estimation algorithm of Picasso Localize (parameters: baseline = 178 photons, sensitivity = 15.5 photons, quantum efficiency = 0.97, box side length = 7 px, min net gradient of 800–2200 photons). Localizations were drift corrected in Picasso render using redundant cross-correlation with a segmentation value of 500 frames. To remove signal from out-of-focus planes, localizations were filtered by their standard deviation (up to a magnitude of 324 nm) and by their localization precision (up to a magnitude of 108 nm). Localizations were linked with a radius of 74.3 nm, which is four times the localization precision of nearest neighbor-based analysis (NeNA) ^33^ and with a maximum dark time of 400 ms ^34^. Localizations were clustered with the density-based spatial clustering of applications with noise (DBSCAN) algorithm ^35^. The radius was set to 74.3 nm and the minimum density to 7 localizations.

### Correlative transmission electron microscopy

Following dSTORM imaging, montaged transmission electron microscopy (TEM) images of the grids were acquired using a 300 kV TEM (Tecnai F30 G2; Thermo Fisher Scientific Inc.) equipped with a charge-coupled device (CCD) camera (UltraScan 4000; Gatan Inc., Pleasanton, CA, USA). Image montages were acquired automatically using SerialEM 3.6.1 ^36^ at a nominal defocus of –5 µm and a total dose of 5.1 e^-^ Å^-2^. The calibrated pixel size was 10.22 Å (×12,000 magnification).

The montaged TEM images were pre-processed by custom MATLAB scripts (MATLAB 2021b, 2021, The MathWorks, Natick, MA, USA). To generate single overview TEM images, we blended montages using the software package IMOD v4.11.7 ^37^. The registration between light and electron microscopy data was performed by selecting control points on both widefield and TEM images and computing an affine spatial transformation using MATLAB (MATLAB 2021b, 2021, The MathWorks). Accuracy of the image registration was compared by overlaying mask and images, respectively. Due to distortions of the sample surface by the preparation, we omitted from our analysis any areas of the sample that showed a high degree of warping.

### Binary mask generation and cluster quantification

The luminal capillary, luminal urinary space, and glomerular filtration barrier were segmented in the EM images using Fiji to create binary masks ^38^. The masks were transformed to fit the rendered image of the clusters, as described above. Cluster densities inside the three segmentations were determined by counting the number of cluster centers inside the respective segmented areas and dividing the counts by the area of the segmented mask with a custom script. The mean and standard deviations were calculated. Paired t-tests were applied to compare the means of the 28 segments in four field of views. All distributions were significantly drawn from a normally distributed population, as determined by a Kolmogorov-Smirnov test (α = 0.05). Levels of significance were classified as p > 0.05, no significant difference (n.s.); p < 0.05, significant difference (*); p < 0.01, very significant difference (**); and p < 0.001, highly significant difference (***). The preparation pipeline is shown in Extended Data Figure 1.

### Isolation of mouse glomeruli

Isolation of glomeruli from the kidneys of wild-type mouse strain C57BL/6J was performed according to an adapted isolation protocol ^39^. All steps were performed at 4°C or on ice. In brief, the kidneys from one mouse were harvested and minced into small pieces (∼1 mm^3^) in Hanks’ Balanced Salt Solution (HBSS, #2323615, Gibco, Thermo Fisher Scientific Inc., Waltham, MA, USA). Minced tissue pieces were gently pressed with a wooden tongue depressor (Glaswarenfabrik Karl Hecht, Sondheim vor der Rhön, Germany), first through a pre-wetted 150 µm cell strainer and then, after extensive washing with HBSS, through a 100 µm cell strainer (PluriStrainer, PluriSelect Life Sciences, Leipzig, Germany). The flow-through was washed again with fresh cold HBSS and pressed through a 40 µm cell strainer. The sample retained on the surface of the strainer was rinsed into a centrifuge tube and centrifuged at 115 *x g* for 5 min at 4 °C. The pellet was resuspended in 100 µL CellBrite Steady Membrane Stain 550 (#30107, Biotium, Fremont, CA, USA) in HBSS (1:1000) and incubated for 25 min, followed by vitrification by high-pressure freezing.

### Sample vitrification by high-pressure freezing

Vitrification of the sample was based on a modified waffle preparation protocol ^40^. Planchets (type B, Wohlwend, Sennwald, Switzerland) were coated using a cetylpalmitate-15 solution (1% w/v in diethyl ether) prior to freezing. Formvar-coated grids (copper/palladium, 100 mesh parallel bar with single bar; G2019D, Plano, Wetzlar, Germany) were placed on the flat side of the planchet, and 3 µL of isolated glomeruli and tubular fragments that had been mixed with an equal amount of Ficoll PM 400 (40% v/v in HBSS) (Sigma-Aldrich, St. Louis, Missouri, US) were applied before the sandwich was completed by adding a second planchet (with the flat side facing the grid). The sample was immediately high-pressure frozen using an HPM-010 (Abra Fluid, Widnau, Switzerland).

### Cryogenic confocal laser scanning microscopy

The grids were imaged at -195°C by confocal laser scanning microscopy (cryo-CLSM) (CMS196 cryo-stage, Linkam, Salfords, United Kingdom; LSM700, Carl Zeiss, Jena, Germany). Optical configurations were adjusted to capture the CellBrite Steady 550 Membrane Stain (#30107, Biotium), and the reflection from the grids in the far-red channel with excitation wavelengths of 555 and 639 nm, respectively. Images were acquired with 5x/NA 0.16- and 20x/NA 0.4 objectives. Montage maps were generated manually using the 20x images and stitched in Fiji v1.51 using the Pair-wise stitching plug-in ^41^. Data acquisition was performed with Zeiss ZEN 2009 (blue) v2.1

### Cryo-focused ion beam milling

The high-pressure-frozen EM grids were clipped into cryo-focused ion beam (FIB) autogrids (#1205101, Thermo Fisher Scientific Inc.) and loaded into a pre-tilted EM grid holder with shutter and additional cold trap under liquid nitrogen using an EM vacuum cryo-transfersystem VCT500 loading station (Leica Microsystems, Wetzlar, Germany). The sample holder was taken up by the EM VCT500 transfer shuttle and transferred into a ACE600 high vacuum sputter coater (Leica Microsystems) for platinum coating (8–10 nm). Via a VCT dock (Leica Microsystems), the sample holder was subsequently transferred into a FIB scanning electron microscopy (SEM) dual-beam instrument (Helios 600i Nanolab, Thermo Fisher Scientific Inc.) equipped with a band-cooled cryo-stage equilibrated at -157°C (Leica Microsystems). The EM grids were imaged using SEM (3 kV, 0.69 nA) and FIB (gallium ion source, 30 kV, 33 pA). An overview SEM image was acquired at 90° incident angle, then an organometallic platinum layer of a few micrometers thickness was deposited gas injection system (GIS) valve open for 5–15 s with inserted GIS needle). Regions of interest were identified by correlating the previous acquired cryo-CLSM with the SEM images using Bigwarp/Fiji. Localized positions were milled by utilizing the specific stress-relief gap for waffled grids as described previously ^40^. The net incident angle varied between 33° and 38°. The following FIB currents were used: 9 nA (trenching), 2.5 nA stepwise down to 83 pA (thinning), 240 pA (notch milling), and 33 pA (polishing). During thinning steps, a second platinum sputtering was done (8–10 nm), followed by GIS deposition of an organometallic platinum layer that was a few micrometers thick.

### Cryogenic transmission electron microscopy imaging and cryo-electron tomography

Cryo-FIB milled lamellae were imaged using a Titan Krios cryogenic transmission electron microscope (cryo-TEM; Thermo Fisher Scientific Inc.) operating at 300 kV in nanoprobe EFTEM mode, equipped with an X-FEG field emission-gun, a GIF Quantum post-column energy filter operating in zero-loss mode and a K2 Summit direct electron detector (Gatan Inc., Pleasanton, CA, USA). Low-magnification images of the lamellae were acquired at the lamella pre-tilt angle induced by FIB milling (pixel size 1.256 nm, –50 µm defocus) with a total electron dose of 0.5 e^−^ Å^-2^ per image using SerialEM v4.1^36^.

Tilt series were recorded at a nominal magnification of ×33,000 (2.18 Å per pixel) in super resolution and dose fractionation modes. The cumulative total dose per tomogram was 150 e^−^ Å^-2^, and the tilt series covered an angular range from −66° to +66° in reference to the lamella pre-tilt, with an angular increment of 1.5° and a defocus set from –7 to –9 µm (Extended Data Table 1). The preparation pipeline is shown in Extended Data Figure 2.

### Cryo-ET image processing

The movie stacks were aligned to correct for beam-induced movement using Motioncor2 v1.6.3 ^42^, and the tilt series were subsequently assembled and aligned using IMOD v4.11.7 patch tracking ^37^. The contrast transfer function (CTF) was estimated using CTFFIND4 v4.1.13 ^43^.

Tomograms for visualization purposes were reconstructed by weighted back-projection using a ramp-filter at a pixel size of 1.747 nm. To enhance the contrast, the Wiener-like tom_deconv deconvolution filter (Tegunov, https://github.com/dtegunov/tom_deconv) was applied and used as input for cryoCARE ^44^ denoising of the tomograms.

Sub-tomograms, which were subsequently subjected to sub-tomogram averaging, were reconstructed by weighted back-projection with exact filter and three-dimensional CTF correction ^45^ at a pixel size of 0.874 nm and 0.437 nm, respectively. From several tilt series, six cryo-electron tomograms with 43 segments of the slit diaphragm (SD) were selected for further analysis, since those were the thinnest and displayed the best alignment.

To measure the distance between two adjacent podocyte membranes and the periodicity of the densities forming the SD by cross-correlation, we used custom MATLAB scripts (MATLAB 2023a, 2023; The MathWorks, Natick, MA, USA).

### Sub-tomogram averaging

For a more precise view of the fishnet arrangement of the SD proteins, sub-tomogram averaging (STA) was performed. Sub-tomograms were pre-oriented such that the top-view direction was aligned with the z-axis and the membrane-view direction was aligned with the x-axis, using the ArtiaX ^46^ tool implemented in ChimeraX v1.6 ^47,48^. An initial model was obtained by averaging 191 pre-oriented sub-tomograms (pixel size of 0.874 nm) with the STA routine of Artiatomi (https://github.com/uermel/Artiatomi) with custom MATLAB scripts. The sub-tomogram positions were refined using a mask containing one membrane of the SD. After initial alignment on the membranes, sub-tomograms diverging from the membrane orientation visible in the tomograms were excluded, yielding the final set of 134 sub-tomograms (pixel size of 0.437 nm). The sub-tomogram alignment was refined using a second mask, excluding the membranes and containing only the SD densities, yielding the final map at a global resolution of 4.7nm (Extended Data Figure 3).

### Segmentation and rendering surface models

Segmentations were manually produced using the modelling tools of ArtiaX ^46^ in ChimeraX v1.6 ^47,48^, and the smoothness of the segmentations was improved using mean curvature motion ^49^ (https://github.com/FrangakisLab/mcm-cryoet).

### Fitting the Nephrin-Neph1 heterodimers to the cryo-ET density maps

The molecular model of Nephrin-Neph1 heterodimers was generated from predictions available from the AlphaFold Protein Structure Database ^18,19^ (IDs: AF-Q9QZS7-F1, AF-Q80W68-F1), since no full-length structure is available to date. The intracellular, transmembrane and signal peptide sections of individual Nephrin and Neph1 AlphaFold predictions were removed and then adjusted according to the density using Coot ^50^ (residues 36-1079 and 48-531 were used for Nephrin and Neph1, respectively). Nephrin and Neph1 were then assembled into a heterodimer based on the crystal structure of the SYG-2-SYG-1 heterodimer (ID: 4OFY, ^13^) by aligning the Nephrin-Ig1 and Neph1-Ig1 domains onto the SYG-2-Ig1 and SYG-1-Ig1 domains, respectively, using ChimeraX. The aligned heterodimer was then rigid-body fitted according to the cryo-ET density map using Coot ^50^. Initially, six heterodimers were rigidly fitted into the cryo-ET density using ChimeraX. Next, the heterodimer model, displayed in as a topview, was built by creating a two-dimensional 2 x 4 lattice of heterodimers. Four heterodimers were oriented along the x-axis (cartesian coordinates, the longest axis of dimers faces towards the membranes), propagating according to their estimated periodicity based on the cryo-ET density map. Another four heterodimers were oriented on top of the previous heterodimers, and were subsequently turned by 180° around the normal vector to the top view, followed by a shift of half their periodicity, using ChimeraX. Finally, to optimize the fit of the heterodimer lattice into the symmetrized cryo-ET density with molecular-dynamics flexible fitting using the ChimeraX tool ISOLDE ^20^, we applied distance restraints for each Ig domain but not for the linker regions between them. Additional distance restraints were applied to amino acids in the Nephrin-Neph1 interface that are assumed to form hydrogen bonds as in the SYG-2-SYG-1 interface (ID: 4OFY), keeping the interface intact during fitting (Extended Data Figure 4).

## References

**Extended Data Table 1:** Cryo-ET data collection and processing.

|  |  |
| --- | --- |
| **Data collection** |  |
| **Microscope** | FEI Titan Krios **G2** |
| **Detector** | Gatan K2 Summit |
| **Acquisition Software** | SerialEM v4.1 |
| **Magnification** | 33,000x |
| **Voltage (kV)** | 300 |
| **Total Electron exposure (e<sup>-</sup> Å<sup>-2</sup>)** | 150 |
| **Defocus range (µm)** | -7 to -9 |
| **Number of frames** | 30 |
| **Images per Tiltseries** | 63 |
| **Increment step** | 1.5 |
| **Pixel size (Å) (super-resolution)** | 2.18 |
| **Processing** |  |
| **Tilt series acquired** | 47 |
| **Tiltseries used** | 6 |
| **Initial particle number** | 191 |
| **Final particles** | 134 |
| **Map Resolution (Å)** | 47.26 |
| **FSC threshold** | 0.143 |
| **Symmetry imposed** | C1 |
| **Motion correction** | Motioncor2 v1.6.3 |
| **Tomographic alignment** | IMOD v4.11.7 patch tracking |
| **CTF estimation** | CTFFIND4 v4.1.13 |
| **Tomographic reconstruction** | Artiatomi |

**Extended Data Figure 1:**
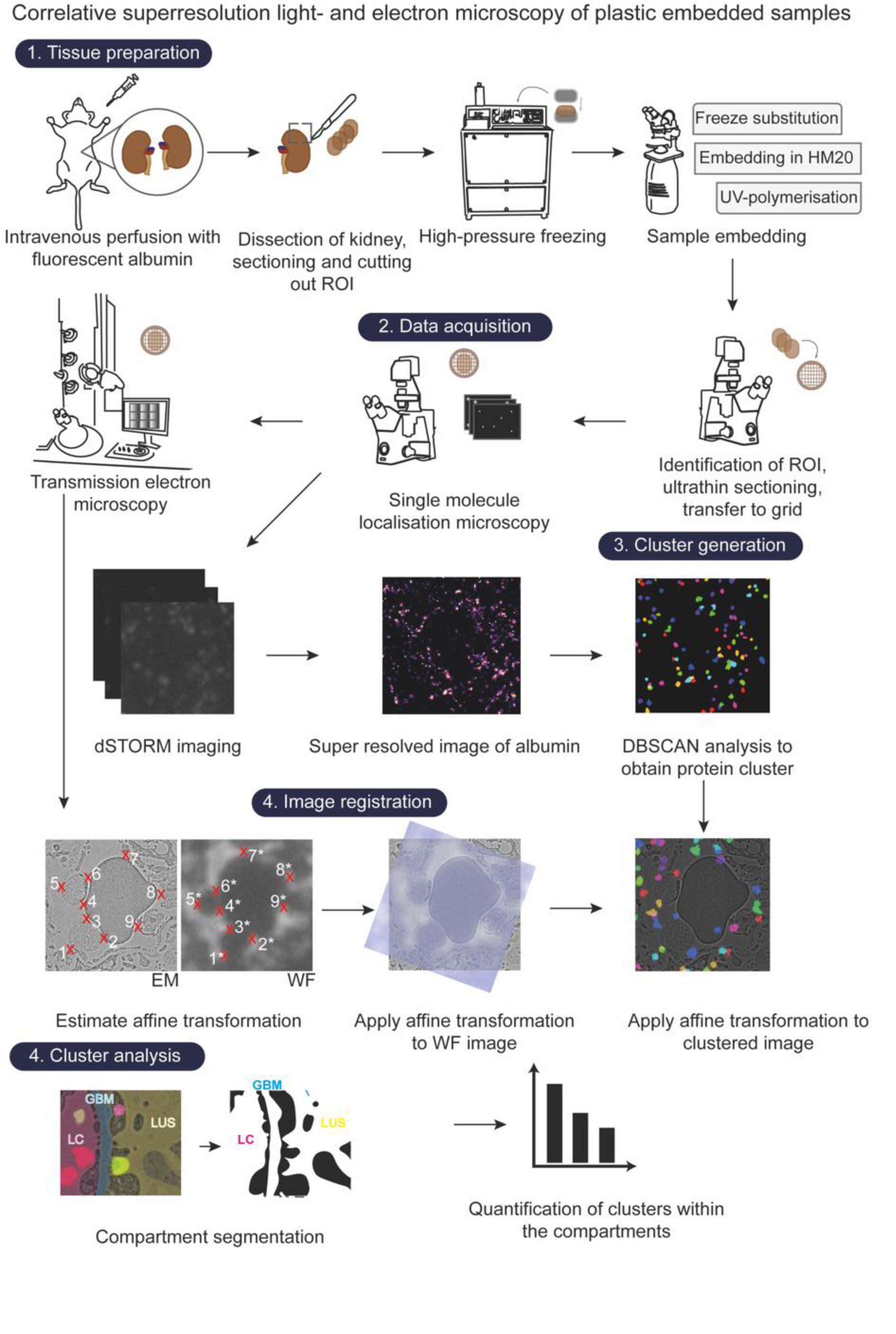
Preparation and processing pipeline for plastic-embedded kidney tissue. Fresh biopsies of mouse kidneys are prepared for high-pressure freezing, followed by freeze substitution and plastic embedding. After thin sectioning and transfer onto electron microscopy (EM) grids, the samples are imaged by direct stochastic optical reconstruction microscopy (dSTORM), followed by transmission electron microscopy. DBSCAN analysis is performed to identify albumin clusters. The EM images are then registered with widefield images of the specimen. This transformation is also applied to the cluster analysis. In a last step, binary masks of the different compartments – the luminal capillaries (LC), glomerular basement membrane (GBM) and lumen of the urinary space (LUS) – are generated and used for cluster quantification.

**Extended Data Figure 2:**
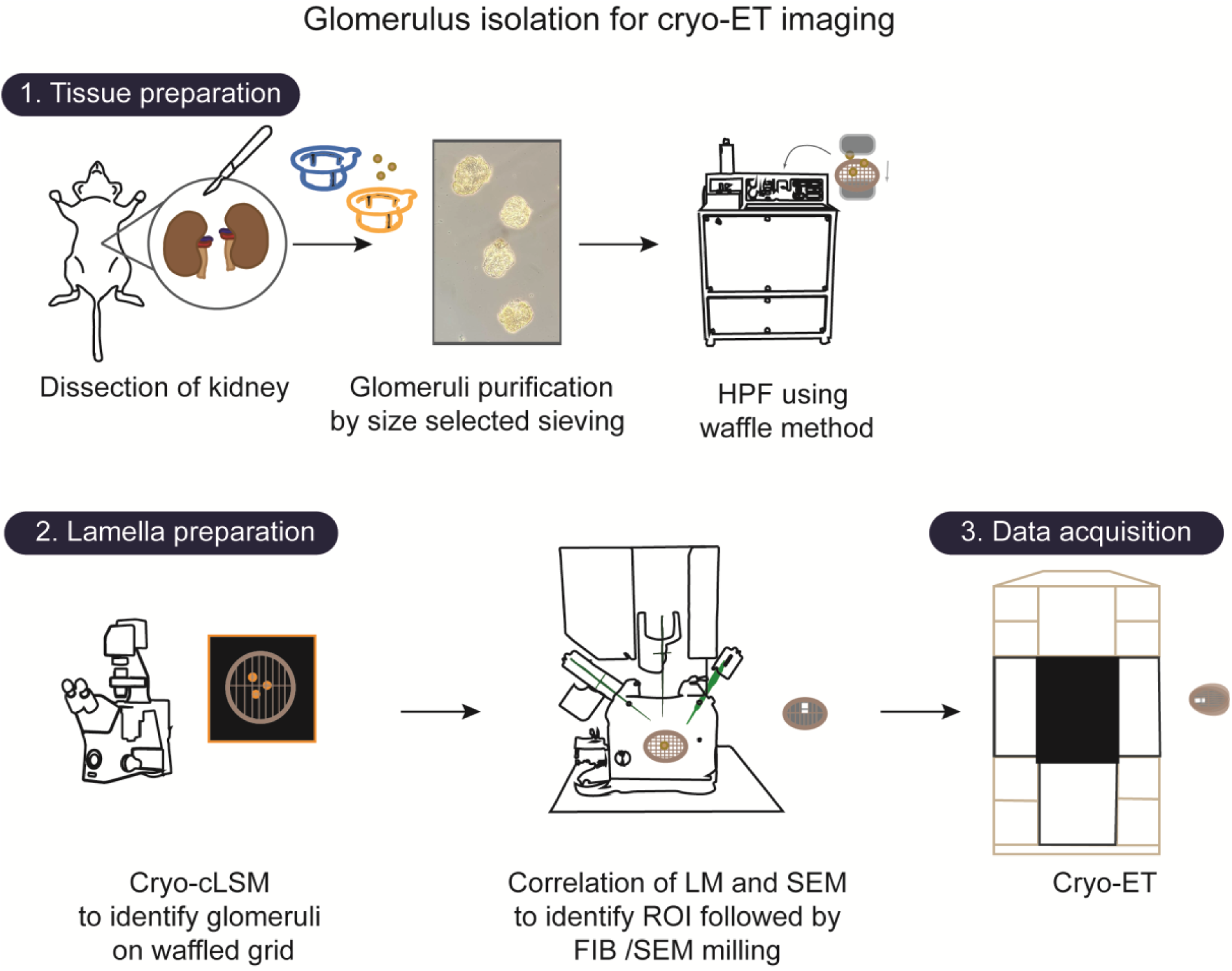
Glomeruli preparation workflow. Glomeruli are isolated from mouse tissue by size-selective sieving and subjected to high-pressure freezing. An overview of the sample is then created using cryo-confocal laser scanning microscopy. The overview is used for identifying regions of interest for the lamellae preparation by means of focused-ion beam scanning electron microscopy milling. Finally, lamellae are imaged using cryo-electron tomography.

**Extended Data Figure 3:**
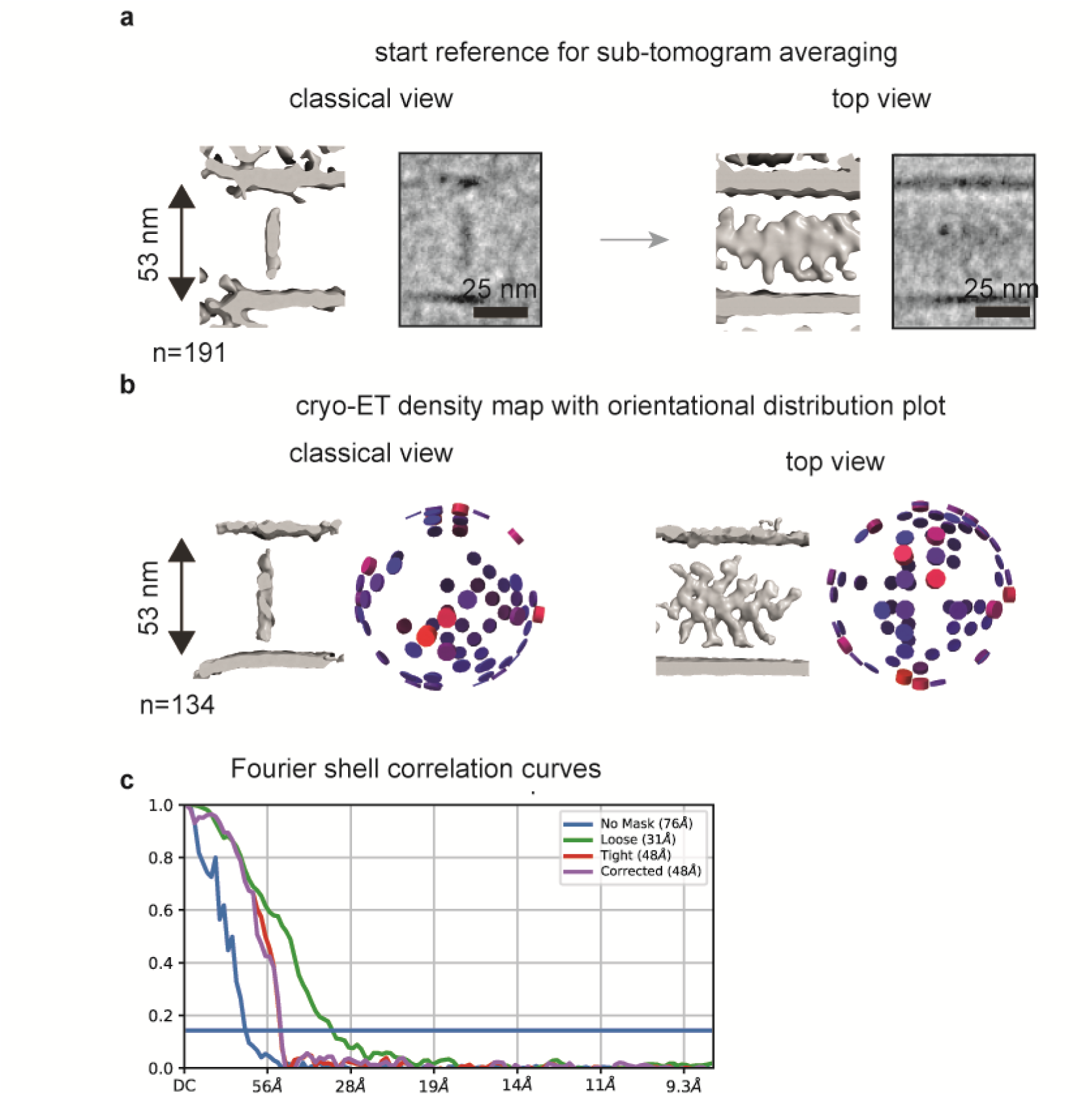
Sub-tomogram averaging and resolution. **a,** Starting reference for sub-tomogram averaging. A homogenous density can be seen in the center of the SD without a discernible structure. **b,** Sub-tomogram average after refinement and removal of outliers. A period structure emerges. Next to the average the rotational position of each average is shown, demonstrating that projections from all views are present. **c,** Gold standard Fourier shell correlation between half-map sub-tomogram averages.

**Extended Data Figure 4:**
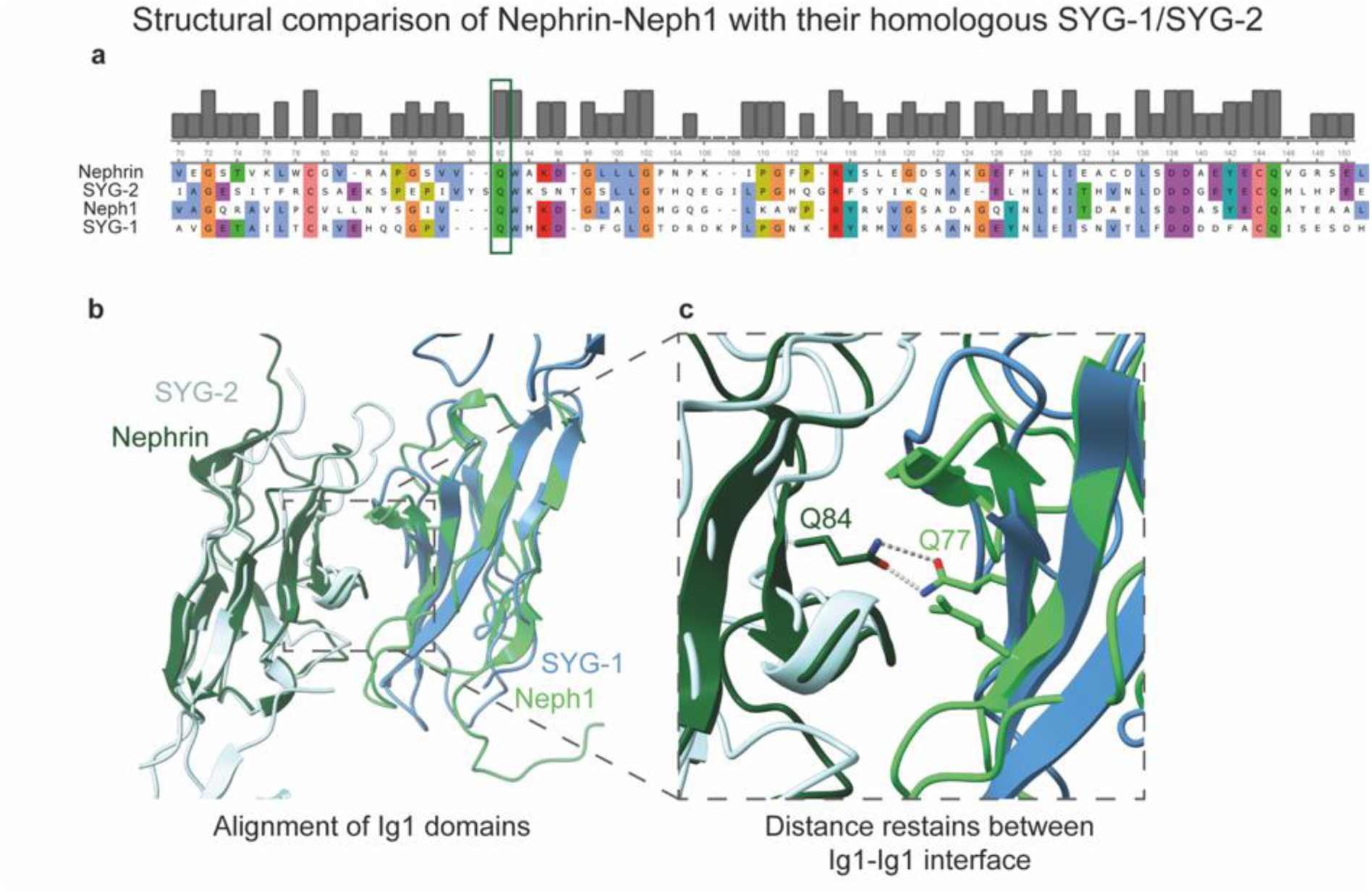
Structural comparison of Ig1 domains of Nephrin and Neph1 with their respective homologs, SYG-2 and SYG-1. **a,** Multisequence alignment of mouse Nephrin, Neph1, SYG-2 and SYG-1 ^51^ ^52^. The sequences are colored by matching amino acids. The conserved glutamine (Q) is highlighted by a green box. **b,** Alignment of the predicted Ig1 domains of Nephrin and Neph1 (AlphaFold Protein Structure Database, IDs: AF-Q9QZS7-F1, AF-Q80W68-F1) ^18^ ^19^ onto the heterodimer structure of SYG-2 and SYG-1 ^13^ (PDB: 4OFY). **c,** For the molecular dynamics flexible fitting, Q84 and Q77 in the Ig1 domains of Nephrin and Neph1, respectively, are distance restrained because the conserved glutamines are mainly responsible for the heterodimerization interface in the SYG-2-SYG-1 heterodimer ^13^.

**Extended Data Figure 5:**
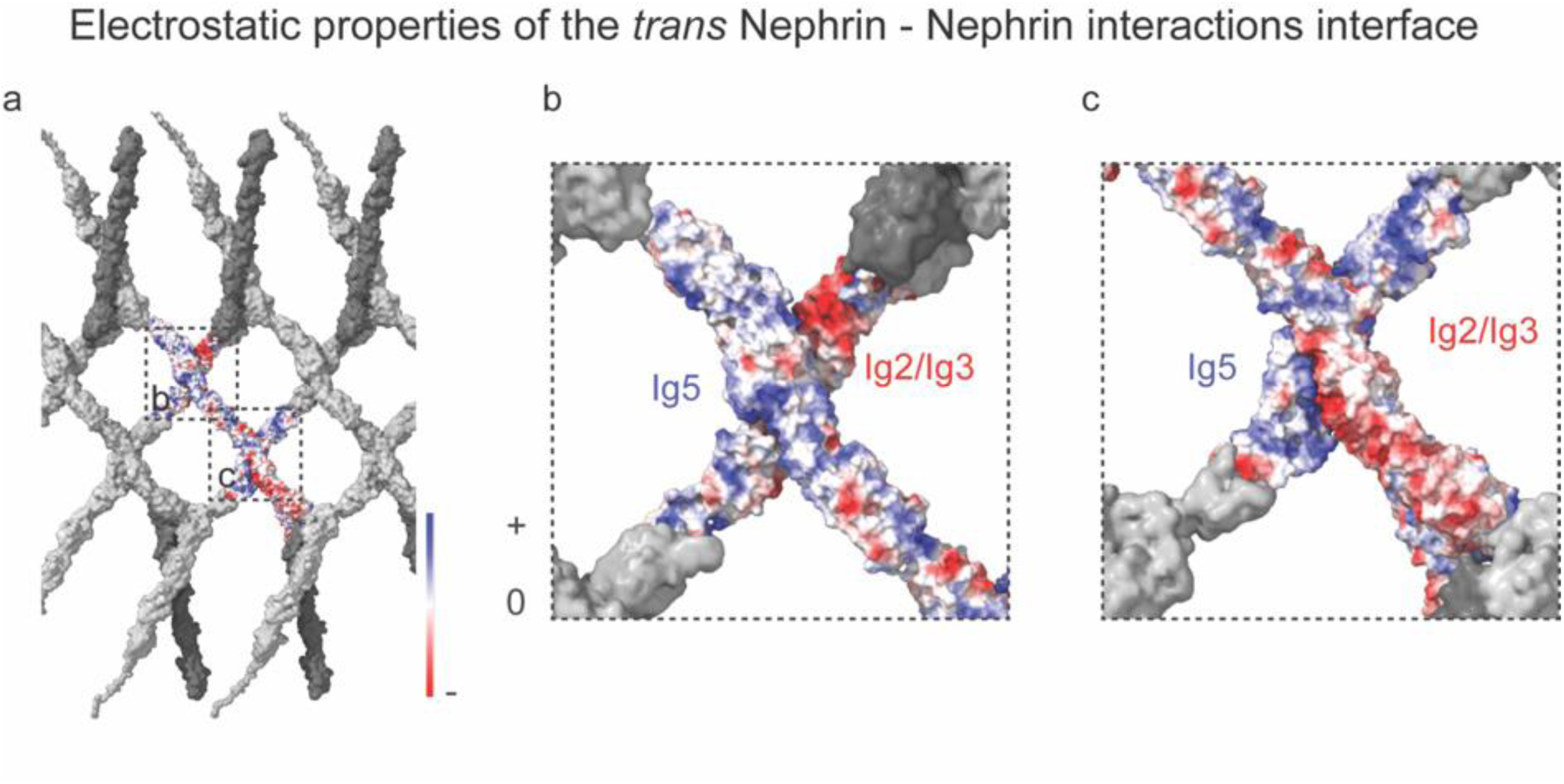
In the model, the electrostatic properties of Nephrin-Ig5 and Nephrin-Ig2/Ig3 are responsible for the homophilic Nephrin–Nephrin *trans* interface. **a,** Top view of the SD model with Nephrin color-coded in electrostatic charge colors. **b,** A magnified view of Nephrin-Ig2/Ig3 and Nephrin-Ig5 from the top view. **c,** The same interface from the bottom view. The region between Nephrin-Ig2 and Nephrin-Ig3 (shown in dark blue in Figure 4c) has negatively charged residues. Nephrin-Ig5 (shown in grey in Figure 4c) is characterized by positive charges.

**Extended Data Figure 6:**
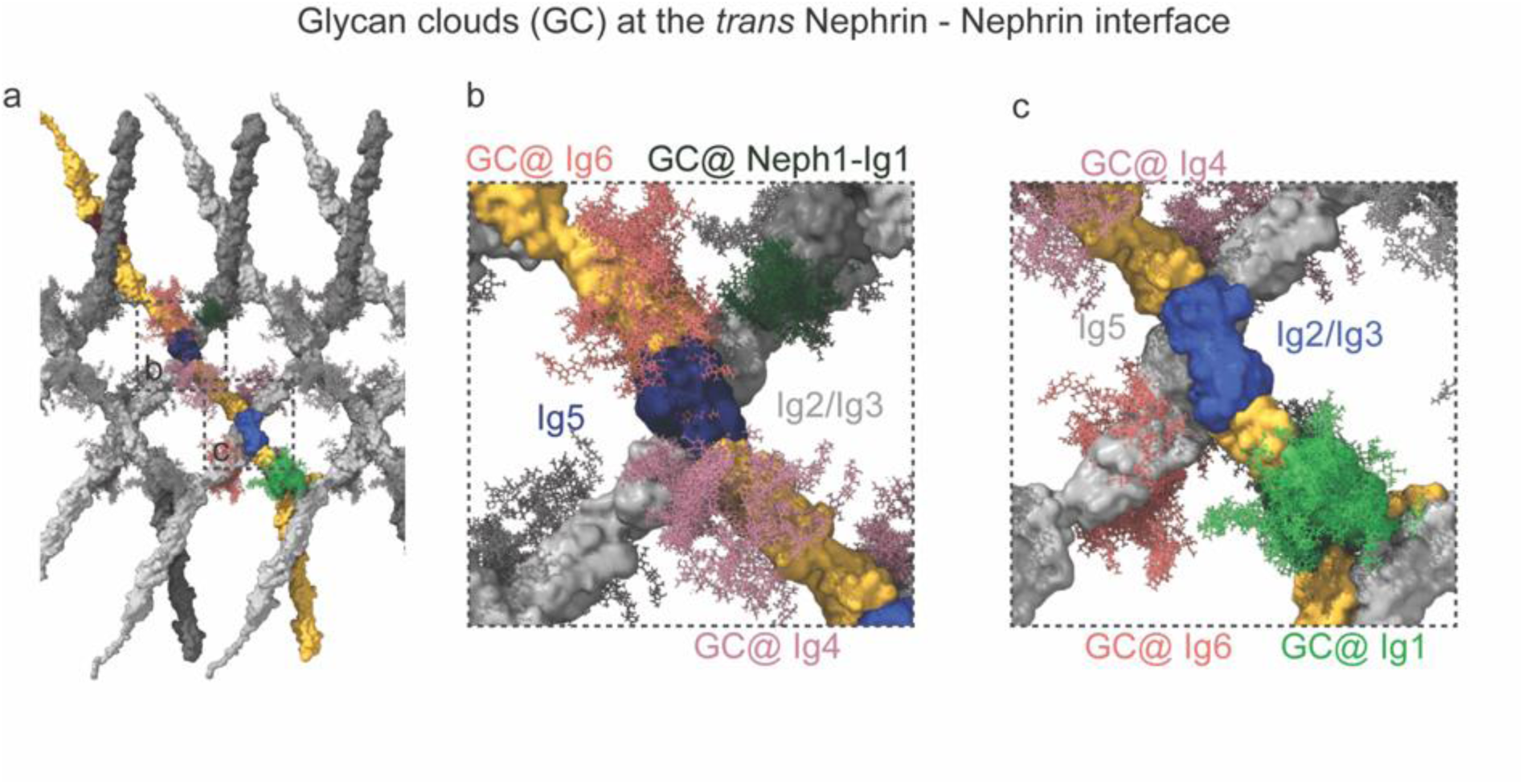
Glycans shield Nephrin Ig domains that are not involved in interactions. **a,** Top view of the SD with the glycans placed on the glycosylation sites of the heterodimer. **b,** Magnified top view of the interface highlighted with “b” in (a) displaying the glycans that leave the interaction domain unshielded. **c,** Magnified top view of the interface highlighted with “c” in (a) displaying the glycans. Nephrin-Ig4 features two glycosylated residues, whereas Ig6 has three. On the side opposite the interface, glycans probably prevent interactions for both Ig4 and Ig6 domains. In addition, the glycans at the tip of the Neph1-Ig1 domain (c), which is oriented toward the Nephrin-Ig2 domain, further constrain the interaction site.

**Extended data movie 1**

Movie of the tilt-series used for the reconstruction of the tomogram shown in Figure 2.

**Extended data movie 2**

Movie of the isosurface visualization shown in Figure 2e.

**Extended data movie 3**

Movie of the isosurface visualization shown Figure 3d.

**Extended data movie 4**

Movie of the model shown in Figure 4.

**Extended data movie 5**

Movie of the model shown in Figure 4 together with the predicted glycans. The glycan cover a significant portion of the structure.

